# Predicting protein variant properties with electrostatic representations

**DOI:** 10.1101/2025.08.11.669649

**Authors:** Floris J. van der Flier, Aalt D.J. van Dijk, Dick de Ridder, Henning Redestig

## Abstract

Does evolution capture the full functional potential of proteins, or is this potential restricted by selective pressures? If the former is true, providing variant effect prediction (VEP) models with evolutionary derived representations should be sufficient to guide the optimization of proteins. In the latter scenario, however, VEP models require different sources of information. In this work, we explore whether physics-based representations of protein variants benefit the performance of VEP models. More specifically, we explore electrostatic representations obtained from solving the Poisson-Boltzmann equation as novel features to fit VEP models to deep mutational scanning (DMS) data. We contrast and combine these representations with those derived from evolutionary models. To this end, we perform a range of experiments: benchmarking, ensembling with evolutionary models, accounting for assay conditions, and extrapolating to new screening data. Though our model displays significant predictive capacity, we find no instance where it provides a better alternative over existing evolutionary models, suggesting that electrostatic representations derived by our methods do not capture extra information compared to evolutionary representations.

## 1. Introduction

### 1.1 Background

Directed evolution is the artificial process of iteratively modifying proteins to improve a set of desired properties. This process can be guided by machine learning models that make predictions about the effect of mutations; a class of models commonly referred to as variant effect prediction (VEP) models. The rate at which improved variants are identified through this process is a bottleneck that restricts the efficiency of protein engineering development cycles. VEP models are therefore a primary target for improvement, which is reflected by the large body of research that introduces new prediction strategies. However, benchmarking efforts indicate that existing models have significant room for improvement. Specifically, the ProteinGym includes a large collection of DMS (deep mutational scanning) datasets with predefined cross-validation (CV) schemes (1). Two CV schemes (described in detail in 2.2) are designed to measure the extrapolative qualities of models by creating test folds containing mutations at positions not mutated in the training data. Models considered state of the art perform significantly worse on these position-exclusive CV schemes when compared to a random CV scheme (2), signaling that there are still performance gains to be made in particular when extrapolating.

The evidence provided by the benchmarking efforts of the authors of the ProteinGym demonstrates that evolutionary models yield the best performance across all VEP datasets. By training on a large collection of naturally occurring protein sequences, evolutionary models are able to approximate an evolutionary distribution of sequences under which their structure and natural function are preserved. While having proved to be useful for various prediction tasks, the information contained in such distributions may be limited in some contexts. In protein engineering, we are interested in the distribution under which desired attributes *for a specific application* are preserved. While the evolutionary and applications-specific distributions are different, they are nonetheless related, at the bare minimum by sharing statistical constraints that preserve primary properties, such as expressibility, structural stability, and function. However, evolutionary and application-specific distributions begin to diverge significantly when properties of interest are no longer relevant to the fitness of the organism from which it was originally sampled. The usefulness of predicting such properties with representations derived from an evolutionary distribution may therefore be limited.

Gingko Bioworks provided evidence of this hypothesis by examining zero-shot scores of protein language models (pLM). These scores are obtained by aggregating predicted likelihoods of each amino acid in a given sequence, and are known to correlate with various protein properties (3). Ginkgo Bioworks showed that enzyme specificity on non-natural targets negatively correlated with zero-shot scores of pLMs (4). Furthermore, while in deep learning, scaling laws of transformers dictate that performance increases with increasing model size, dataset size, and compute, these scaling laws do not carry over to downstream prediction tasks, like VEP, as demonstrated by Li et al. (5). A large pretraining experiment by Gingkgo Bioworks empirically corroborated this finding, demonstrating that almost tripling pretraining size by expanding UniRef sequences with metagenomic sequences had a negligible effect on VEP performance (4).

While currently offering top of the line performance, evolutionary models thus have serious limitations that impair their applicability to VEP. Featurization methods that are not constrained by the evolutionary landscape of proteins are therefore a logical venue to explore. Complementary to the data-driven approach, which involves pretraining on extensive samples of evolutionary data, is the first-principles approach – constructing molecular models of protein variants. While fully accurate physical representations of protein variants under assay conditions may not be feasible, there are methods available to generate reasonable approximations of such models.

Other than providing a complementary source of information with respect to evolutionary methods, explicit physical representations of protein variants may facilitate learning by directly providing models with features that have near linear statistical relationships with properties of interest. Such a relationship is exemplified in a patent published by Cascao-Pereira et al., where stain removal activity of a protease in laundry application was shown to be strongly influenced by the enzyme’s charge (6). Lastly, another attractive attribute of physical representations is that – in contrast to current sequence-based representations – they are contextually flexible by taking into account physical conditions like the temperature, ionic strength, pH, etc.

Physical features of proteins have previously been explored to model the effect of protein mutations. (7) and (8) represented protein variants using voxel representations with channels corresponding to different biophysical quantities and used a 3D CNN to make protein binding affinity and stability predictions, respectively. These methods have the drawback of assigning physical quantities on a per atom basis, and therefore do not act on a realistic representation of the physics of the system –*i*.*e*., the volume that harbors the protein – which is ultimately what provides a physically grounded explanation of downstream assayed properties of protein variants.

### 1.2 Calculating electrostatics with solvent models

The charge distribution and resulting electrostatic potential describe the physics of the protein variant to a more detailed extent than per-atom representations of physical quantities. They are not confined to individual species of the system, like individual atoms or residues, but instead capture how the distribution of charge within a molecule gives rise to long-range electrostatic effects. The electrostatic potential represents the potential energy a hypothetical unit charge would experience as a result of surrounding charges, including both fixed charges on the protein and redistributed charges in the solvent. It provides insight into both the local physical properties of a protein, like reactivity of the active site and binding potential, as well as global properties, like hydrogen bonding networks governing protein stability (9). It also induces the spatial arrangement of solvent charge density – the distribution of ions –, which plays a significant role in solvation, ion screening, and long-range interactions (10; 11).

Electrostatic quantities like electrostatic potential and charge density can be modeled with explicit or implicit solvent methods. Molecular dynamics is an example of the former, capturing the interactions of all atomic species in the solvent. While offering great accuracy arising from detailed solute-solvent interactions, such methods sacrifice speed and require extensive computational resources due to the large number of degrees of freedom of all atoms in the system. This holds especially true when dealing with larger molecules such as proteins. Implicit solvent methods avoid dealing with solvent degrees of freedom by treating the solvent as a continuous medium and describing its aggregated influence. There are many such methods available, each relying on a different set of assumptions that can be matched to specific scenarios. Among these methods, the generalized Born model is a popular method that dates back to 1990 and is continuously developed (12). Though this method is computationally lightweight, it is of limited use when long-range interactions and fine-grained resolution are of importance.

Over the last few decades, methods that solve the Poisson-Boltzmann equation have become the gold standard in characterizing electrostatics for larger biomolecular structures (13). When compared to the generalized Born model, these methods provide a global, more detailed depiction of the electrostatics, including the interior of the biomolecule, which is a strong determinant of downstream protein properties. Finite element and finite difference methods are currently the most broadly used solvers within this domain, with the former trading in speed for increased accuracy and fine-grained resolution with respect to the latter.

In this work, we explore the potential of using electrostatics in VEP. We do so by building ChargeNet: a 3D CNN designed to predict the effect of mutations in proteins based on their electrostatic properties generated by a finite difference solver of the Poisson-Boltzmann equation (PBE). We populate voxel channels with the charge distribution, solvent charge density, and electrostatic potential to provide the network with a comprehensive picture of the electrostatics of variants in our dataset. Though these quantities are strongly related, we include all three in our feature representation, since some may be more directly related to certain properties of interest than others, and may therefore facilitate prediction.

We perform a range of experiments that assess the utility of ChargeNet in VEP. We evaluate the salience of simple charge-derived quantities, benchmark ChargeNet against evolutionary models on various extrapolation tasks, assess the flexibility of accounting for assay conditions, and determine whether electrostatic representations can be used to complement evolutionary models through ensembling. We also evaluate ChargeNet in an artificial scenario in which we expect it to outperform evolutionary models as a control experiment. While we ultimately find that ChargeNet does not provide benefit over evolutionary models in terms of in silico prediction performance in the evaluated datasets, we also show that ChargeNet can still achieve reasonable performance on challenging extrapolation tasks, without relying on any form of pretraining.

## 2. Results

### 2.1 Predicting protein variant properties with electrostatic representations

Before describing our experiments in detail, we first describe the components that make up ChargeNet in Figure 1. All experiments in this work investigate the supervised learning capabilities using these components, or a subset thereof. ChargeNet operates on protein sequence-property pairs and consists of a physics module that computes electrostatic representations, and a deep learning module that operates on these representations, finally yielding a prediction of a property of interest.

**Figure 1.**
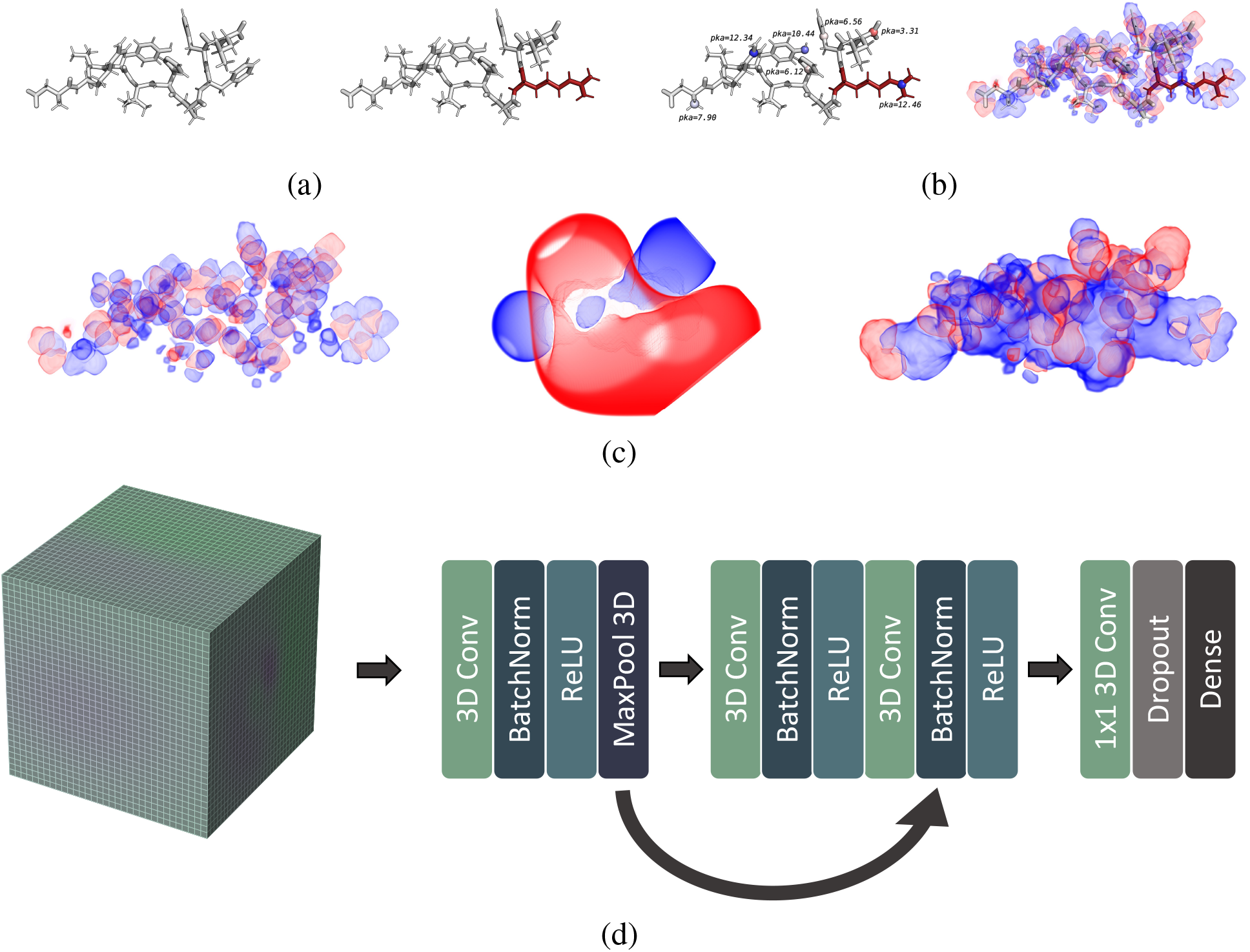
An overview of the steps involved in the ChargeNet pipeline using angiotensin I (PDB ID: 1N9U) for illustrative purposes. We visualized structures and volume renders of electrostatic quantities using PyMOL (14). In all volume renders, red colors correspond to negative values, while blue colors correspond to positive values. (a) FoldX predicts the structure of a variant (right) using a reference structure (left) and the mutation(s) corresponding to the variant sequence (15). The mutation in this example is F8K and is colored in red. (b) PDB2PQR computes the pKa’s of all titratable groups in the structure (left), and then determines the protonation status of those groups given a pH. With this information, PQDB2PQR assigns partial charges and charge radii of the protein variants (right) and stores them in a PQR file. (c) APBS solves the Poisson-Boltzmann equation for the charge assigned structures given a configuration file with solver options and additional specifications of the chemical environment. APBS then outputs the charge distribution already computed by PDB2PQR, the solvent charge density, and the electrostatic potential as voxel grids, here illustrated from left to right. Note that the visualizations here are volume renderings created in PyMOL, shown to facilitate interpretation of the quantities, and do not reflect the computed voxel representations. (d) These three quantities are stored in a 4D tensor, with three channels corresponding to the electrostatic quantities produced by APBS. A small 3D CNN with a ResNet architecture (16) is then trained to predict corresponding screening measurements of those variants.

The first component of the physics module invokes a mutagenesis tool, in this case FoldX (15), to predict structures of protein variants based on a rotamer library and a structure of the reference structure – *i*.*e*., a structure of the unmutated protein sequence. Next, the PQR component assigns the atoms in the predicted protein structures with partial charges (q) and radii (r) using PDB2PQR (17), based on a user specified pH value. PDB2PQR relies on PROPKA3 (18) to predict the pKa values of all titratable groups in the protein – *i*.*e*., parts of the protein that can be (de-)protonated.

We calculated the electrostatic quantities using the finite difference solver implemented in APBS (Adaptive Poisson-Boltzmann Solver), as this ultimately yields the best tradeoff of performance and speed. APBS offers efficient implementations of multiple PBE solvers and is built to be compatible with PDB2PQR, thereby facilitating streamlined computation of electrostatic quantities from structure. The configuration used to perform these computations is described in Methods 4.3.

Finally, the volumetric quantities obtained from APBS are used to populate the channels of a voxel grid, yielding a 3D image of the electrostatics of the protein variants. These voxel grids are then used as inputs to a shallow 3D CNN with a ResNet-like architecture (16), which is trained to predict an assayed property.

### 2.2 The electric dipole moment is a weak predictor of variant effect

We started from a very simple question: does the dipole moment, a vector abstraction of the charge distribution, harbor any predictive capacity in VEP? Should this be the case, it would demonstrate that the orientation of charges in proteins is a salient feature in VEP and that its use is worth investigating further. From a model development perspective, it also provides a baseline performance, informing us of the added value of globally resolved electrostatic representations and feature extraction capabilities of the CNN.

The dipole moment of molecules with non-neutral charge is easily computed by summing coordinate vectors of all atoms, 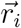 relative to a point of observation 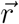, weighted by their charge *q*_*i*_ (19; 20):

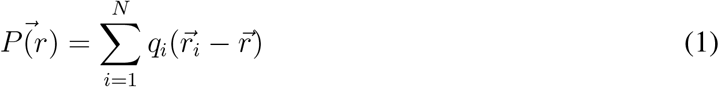

The point of observation customarily corresponds to the center of mass of the molecule in the case of proteins (21). We can find the vector corresponding to the center of mass as:

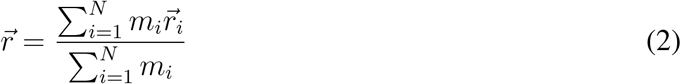

with *m*_*i*_ denoting the atomic mass of atom *i* in the structure. We use the previously described mutagenesis and PQR components of the pipeline to obtain the coordinates and corresponding partial charges of all of our variants, and use a simple regression model to make predictions about the properties of our variants with the three-dimensional dipole moments as inputs. The resulting pipeline is what we shall henceforth refer to as the dipole regressor.

We chose a representative linear and non-linear regression model to make dipole-property predictions, ordinary least squares (OLS) regression, and a second-order polynomial regression model (POLS). The reason for choosing these two models is that a linear model is by definition monotonic and is unable to segment the dipole space into regions of high and low property values, whereas a non-linear model can. We hypothesize that this aligns better with reality, as the relationship with the dipole moment and the associated property values is more likely to exhibit optima rather than a continuously increasing trend. The comparison between the linear and polynomial models should reflect the extent to which this holds for a particular dataset and should tell us if increasing model complexity yields benefit.

We evaluated the dipole regressors with both prediction heads on a curated selection of datasets from the ProteinGym. Further information on data curation is presented in Methods 4.1. We obtained Spearman correlation estimates using K-fold cross-validation (CV) with the contiguous, modulo, and random splitting strategies (1), and we organized these results in Figure 2. The contiguous scheme creates folds of mutated sequence positions spanning contiguous segments of the sequences. *e*.*g*., for a five-fold contiguous CV, fold 1 contains sequences with mutations at the first five positions, fold 2 at positions 6 to 10, etc. The modulo scheme creates folds based on the modulo of the mutated position number of the sequence. *e*.*g*., for a 5-fold modulo CV, sequences with mutations at position 1 are assigned to fold 1, 2 to fold 2, 6 to fold 1, etc.

**Figure 2.**
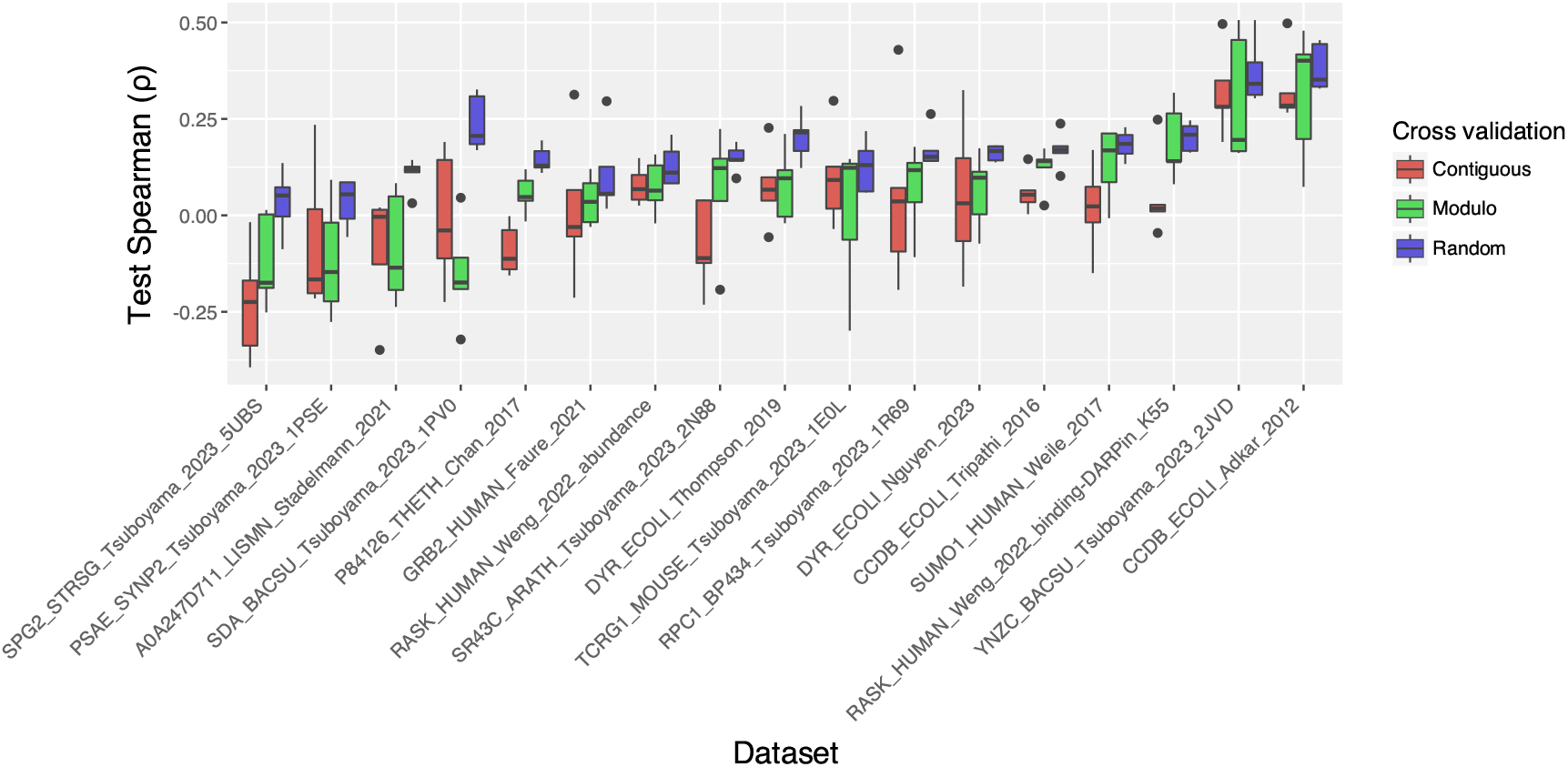
Spearman correlation coefficient scores of the dipole regressor with the POLS prediction head on five test folds using a contiguous, modulo, and random CV strategy for the curated ProteinGym datasets. Correlations are weak overall, but some datasets display a clear relationship between the dipole moments of variants and their assayed properties.

The average and standard deviations of the accuracy of both dipole regressors, along with all splitting strategies and all datasets, are organized in Table S2. The dipole regressor with the POLS prediction head outperforms its OLS counterpart, corroborating our hypothesis about the non-monotonic relationship between dipole moments and screened properties, indicating that there is more to gain from adding model complexity. Despite overall achieving low Spearman correlation, the POLS dipole regressor is weakly predictive of variant effect in some of the datasets, demonstrating that charge-derived representations can act as informative features in VEP models. Median and maximum average Spearman correlations across datasets were 0.16 and 0.38 for random CV, 0.07 and 0.31 for modulo CV, and 0.10 and 0.27 for contiguous splitting. Model performance varied with splitting strategy, showing decreasing correlations and increasing variance from random to modulo to contiguous splits. An attributing factor to this is that the dipole distributions differ significantly between folds for position-based splitting strategies, increasing the difficulty of the prediction task. We determined the difference between the distributions of the magnitude of the dipole moment by calculating the respective Wasserstein distances and displayed these results in Table S1. It is clear from these results that the higher Wasserstein distances by split match the lower Spearman scores of the model, suggesting that domain shift is responsible for decreased performance.

### 2.3 ChargeNet is not competitive with evolutionary models on ProteinGym datasets

Having established that the dipole moment can be used as a feature for VEP, we chose to investigate whether providing a more complex model with globally resolved charge distribution and additional electrostatic features could improve prediction performance to the extent of outperforming VEP models leveraging evolutionary features. We repeated the dipole regressor analysis with ChargeNet to learn to answer this question. We are particularly interested in scoring ChargeNet with the modulo and contiguous CV strategies. Random splitting strategies result in training and testing sets that generally have a large degree of overlap in which positions are mutated. Training models on feature vectors composed solely of indicator values that indicate for each position in the sequence whether it is mutated is often enough to obtain decent Spearman scores (22). Scoring models with random splitting strategies is therefore relatively uninformative for the extrapolative qualities of a model.

We compared ChargeNet with two evolutionarily informed models, the augmented Potts model and the RITA regressor (22) and organized the performance scores aggregated by dataset in Table 1), and for each dataset and CV strategy in Tables S3, S4 and S5. While reaching meaningful performance in both position-based extrapolation CV strategies, ChargeNet is significantly outperformed by the evolutionary models on every dataset and CV strategy, and thus does not prove to be a meaningful alternative to evolutionary models in terms of prediction performance.

**Table 1.** Dataset aggregated averages and standard deviations of Spearman correlation coefficient scores for all CV strategies. Model scores on individual datasets are reported in Tables S3, S4, S5 for the random, modulo, and contiguous CV strategies, respectively.

| CV strategy | Model name |  |  |
| --- | --- | --- | --- |
|  | ChargeNet | Augmented Potts model | RITA regressor |
| Random | $0.59 \pm 0.10$ | $0.73 \pm 0.10$ | **$0.77 \pm 0.11$** |
| Modulo | $0.29 \pm 0.18$ | $0.44 \pm 0.18$ | **$0.52 \pm 0.17$** |
| Contiguous | $0.25 \pm 0.15$ | $0.43 \pm 0.20$ | **$0.46 \pm 0.22$** |

### 2.4 A stacking ensemble of ChargeNet and evolutionary models does not outperform single evolutionary models

Although ChargeNet does not outperform evolutionary models on any of the ProteinGym datasets, it was able to produce meaningful predictions (Spearman > 0.2) for nearly all datasets and CV strategies. Since ChargeNet and the evolutionary models use entirely different feature extraction methods, we were interested in combining their predictions. We reason that ChargeNet predictions might use information not considered in predictions of evolutionary models, and that training a meta learner to yield ensemble predictions might help in finding a combination of these predictions that would result in increased performance.

We examined this through a procedure called stacking (23). We provide a detailed description of our implementation in Methods 4.8. In short, data is divided into five folds, of which three for training, one for validation, and one for testing. The training folds are used to train ensemble members, and the validation folds are used to train a meta learner, which in our case is a simple ridge regression model. The meta learner is trained to predict ground truth values using feature vectors with model predictions as inputs. Since our model is a ridge regression model, the regression coefficients are the weights of the ensemble member predictions, and the bias term offsets weighted predictions.

We compared the test predictions of ensemble members and all possible combinations thereof on our curated set of ProteinGym datasets using the modulo CV strategy (Figure 3). We chose this CV strategy to save on computation, and since it ranks between the random and contiguous CV strategies in terms of difficulty, providing a moderately difficult scenario to evaluate the predictors. It should be noted that the scores reported for the ensemble members in this experiment are different

**Figure 3.**
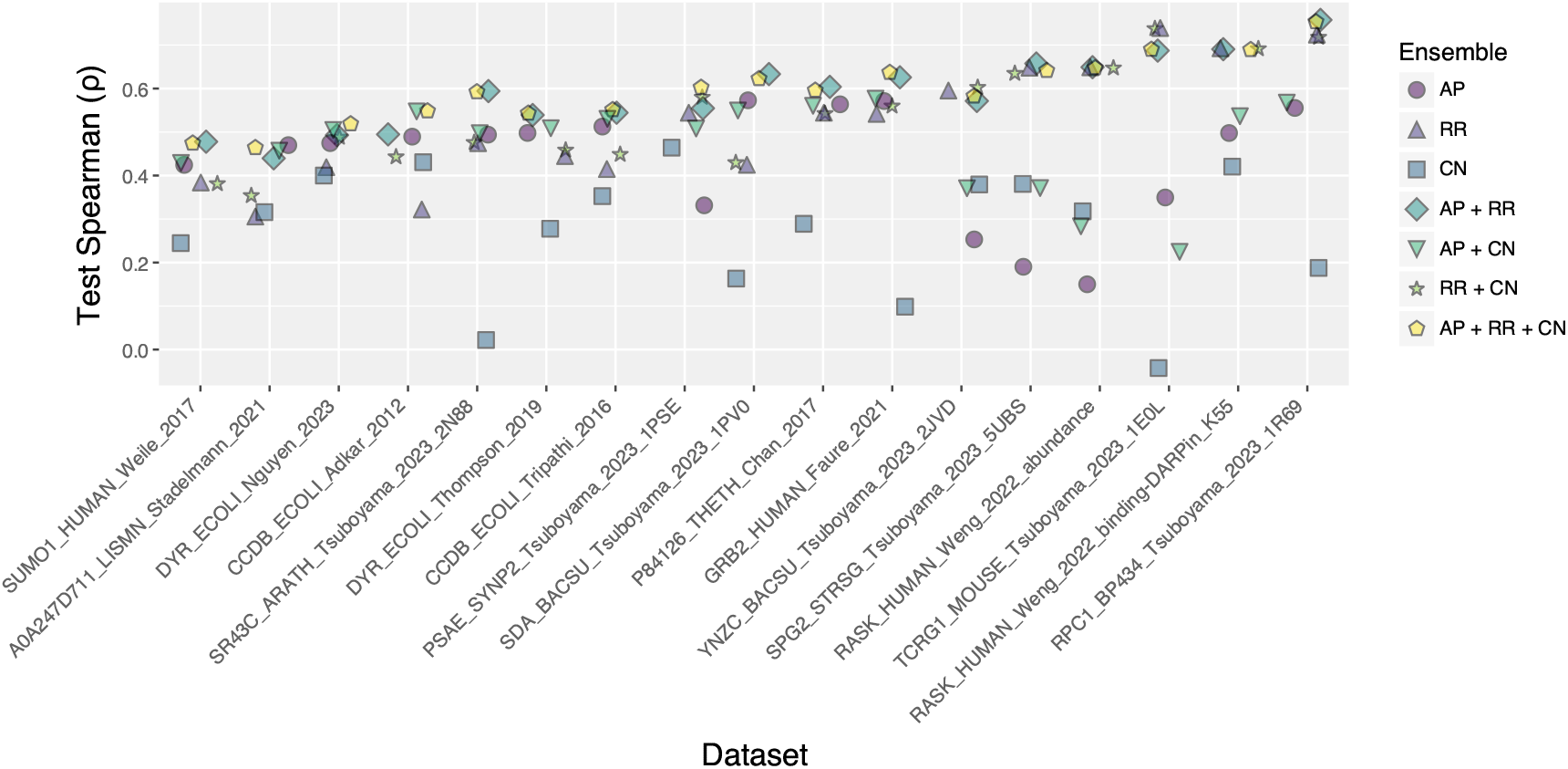
Average Spearman correlation coefficient scores of ensemble and standalone models for the curated Prote-inGym datasets over all five test folds with the modulo CV strategy. Model name abbreviations are: AP: augmented Potts model, RR: RITA regressor, CN: ChargeNet.

from the scores for datasets using the modulo CV strategy in the previous experiment (Table S4), the difference being that we trained on three out of five folds in the former and on four out of five folds in the latter.

While there is marginal benefit to ensembling both evolutionary models, there is no example where adding ChargeNet as an ensemble member to a single evolutionary model increases test performance in terms of Spearman correlation, making the use of ChargeNet as a member in a stacking ensemble ineffective.

### 2.5 ChargeNet is not competitive with evolutionary models in predicting activity of a future screening round of *α*-amylase

In the previous experiments, we investigated extrapolation ability within the scope of a single dataset. However, it is often of more direct interest in protein engineering how well prediction algorithms can predict the outcome of a future screening round. The data necessary to conduct such an experiment has been made available by the Protein Engineering Tournament (24). In this tournament, the authors provided participants with a dataset of *α*-amylase variants and corresponding expression, specific activity, and thermostability measurements that they could use to learn from and generate new variant sequences that would maximize activity. Chase et al. have made the submitted sequences available, which can be used by model developers to probe extrapolation quality to new screening data.

We trained ChargeNet alongside the augmented Potts model and RITA regressor on five random 80% subsets of the training data, *i*.*e*, data made available prior to the Protein Engineering tournament. After each training run, we evaluated the models on all test data – *i*.*e*., variants that were submitted by participants of the Protein Engineering Tournament and screened. In doing so, we obtained five estimates of model performance on the test data, which are 0.24*±* 0.06, 0.42 *±*0.02, and 0.40 *±*0.02 for ChargeNet, the augmented Potts model, and the RITA regressor, respectively. From these results, we must conclude that ChargeNet is also outperformed by the evolutionary models on the task of extrapolating to newly generated screening data.

### 2.6 ChargeNet performance is not affected by the pH under which electrostatics are computed

So far, we have directed our focus to the applicability of ChargeNet as a supervised predictor of variant effect. Having not found any evidence of benefit over evolutionary models, we wanted to investigate a potentially attractive feature of ChargeNet, namely the ability to operate on electrostatic representations that match the screening conditions of the datasets. To this end, we chose two datasets to train ChargeNet at the pH described in the corresponding studies, and contrasted that with ChargeNet trained at pH values close to the extremes, *pH* = 1 and *pH* = 13. We selected the datasets based on two criteria: the pH value of the dataset needed to be close to neutral, and ChargeNet performance with a position based splitting strategy needed to be adequate (Spearman ⪆ 0.3). We reasoned that it would be more difficult to contrast performance scores when performance was either very high (not observed) or very low. We refer to the results presented in Tables S4 and S5 for the latter criterion. With these criteria in mind, we picked the PSAE_SYNP2_Tsuboyama_2023_1PSE and YNZC_BACSU_Tsuboyama_2023_2JVD datasets.

We trained three different models using 5-fold CV with the modulo splitting strategy to obtain performance estimates for three pH values: the lower extreme, 1; the correct pH, 7.4; and the upper extreme, 13. We accounted for pH values in the partial charge assignment by PDB2PQR, and in the PB solving step of APBS. Table 2 reports the Spearman correlation scores of the models trained with this procedure, from which it can be concluded that the model trained on electrostatic representation generated at the correct pH does not outperform the others.

**Table 2.** Averages and standard deviations of Spearman correlation coefficient scores of ChargeNet with three different pH settings (lower extreme, correct, upper extreme) on all five test folds of two selected ProteinGym datasets using the modulo CV strategy.

| pH<br>Dataset | 1.0 | 7.4 | 13.0 |
| --- | --- | --- | --- |
| PSAE_SYNP2_Tsuboyama_2023_1PSE | $0.45 \pm 0.17$ | $0.42 \pm 0.09$ | $0.46 \pm 0.15$ |
| YNZC_BACSU_Tsuboyama_2023_2JVD | $0.44 \pm 0.28$ | $0.39 \pm 0.22$ | $0.31 \pm 0.18$ |

What is also noticeable is that the performance estimates tend to have large standard deviations, complicating comparison. To address this, we repeated this experiment with a “softened” CV strategy, *i*.*e*., such that test folds contain sequences with shorter distances to the training set than in the modulo CV strategy. For this purpose, we created the PIME (position inclusive mutation exclusive) split, which we describe in greater detail in section 4.2. Table 3 displays the scores of the previously outlined experiment using the PIME CV strategy.

**Table 3.** Averages and standard deviations of Spearman correlation coefficient scores of ChargeNet with three different pH settings (lower extreme, correct, upper extreme) on all five test folds of two selected ProteinGym datasets using the PIME CV strategy.

| pH<br>Dataset | 1.0 | 7.4 | 13.0 |
| --- | --- | --- | --- |
| PSAE_SYNP2_Tsuboyama_2023_1PSE | $0.67 \pm 0.05$ | $0.70 \pm 0.04$ | $0.64 \pm 0.07$ |
| YNZC_BACSU_Tsuboyama_2023_2JVD | $0.60 \pm 0.07$ | $0.63 \pm 0.05$ | $0.59 \pm 0.05$ |

In contrast to the scores obtained from the modulo CV strategy, using the PIME CV strategy, the highest scores are obtained with electrostatic representations obtained at correct pH conditions for both datasets. These results suggest that models operating on physical quantities have the ability to account for screening conditions, which is a valuable feature when working with screening data obtained at non-physiological conditions. However, we can not definitively base this conclusion on these findings as the differences in scores are not statistically significant for any of the comparisons (lower extreme–correct, upper extreme–correct), as determined by a t-test (*p* > 0.05).

## 3 Discussion

In this work, we explored the potential of using computed electrostatic representations of protein variants to predict properties. The pipeline implementing this approach, which we named ChargeNet, consists of four different components: 1) a structure prediction component that maps variant sequences together with a reference structure to a variant structure; 2) a partial charge and charge diameter assignment component; 3) a PB equation solver component that creates 3D grids of electrostatic quantities; and 4) a 3D CNN operating on those grids for supervised prediction of protein variant properties. To learn if our model could be used effectively for VEP tasks, we carried out a number of experiments that measured different capabilities of our model.

The capabilities our experiments assessed can roughly be categorized as extrapolation, complementarity with existing models, and adaptability to the data. We assessed extrapolation in an intra-screen manner, by evaluating model performance on protein variants with mutations at positions not mutated in the training set, and also in an inter-screen manner, by evaluating model performance on a later round of screening data of the same enzyme. We investigated the complementarity of ChargeNet with existing models by creating stacked ensembles of ChargeNet predictions and evolutionary model predictions. Lastly, we looked at the ability of ChargeNet to account for the chemical conditions of the screening data by computing electrostatic quantities under correct pH values, and ones that are far away from the correct pH.

For all these experiments, we find that ChargeNet in its current implementation does not offer any benefits over using evolutionary models in supervised prediction of protein variant properties. ChargeNet is always outperformed by either the augmented Potts model, the RITA regressor, or both, and including it in an ensemble with those models does not yield any additional performance. While we found some evidence that ChargeNet performs better when pH is accounted for in the physics module, the differences in performance using incorrect pH values were not statistically significant.

### 3.1 Reflections

Since physics is ultimately what underpins all assayed properties of protein variants, a model operating on representations grounded in physics is an appealing topic to explore. While we were not able to find a scenario in which ChargeNet could provide value, we certainly do not want to discourage others from exploring similar strategies. However, we think it is important to reflect on the potential reasons that existing methods outmatch ChargeNet and to envision different directions for physics-based VEP in order to make meaningful progress in this field.

We hypothesize that there may be a number of reasons why ChargeNet performed suboptimally. We group these problems into four categories:

- **Features may poorly reflect biological reality**. Prediction performance of machine learning models is greatly influenced by the extent to which input features match reality. In the case of ChargeNet, representations generated by the physics module do not reflect the biological context of the proteins influencing them from the moment they are synthesized, such as post-translational modifications or molecular crowding effects, nor do these representations capture the dynamics of the protein. Models like ChargeNet that consider only snapshots of proteins have no way of carrying this information. Sandhu et al. recently showed that protein function is often determined by a small subset of states of the conformational ensemble, and that small shifts in the relative population of these states can significantly impact the function of the protein (25). Lastly, while ChargeNet does have the ability to account for the ionic composition of the solvent, this information can not always be accurately determined and may also be transient.
- **The physics module may provide erroneous results**. Not only are these representations incomplete reflections of biology, their generation is susceptible to multiple sources of error. The physics module consists of four components that are all prone to errors, and their serial application can compound these. This can ultimately lead to large differences in estimates of the electrostatics versus the ground truth. Significant potential errors, listed in the order they may occur within the pipeline, include:
  – FoldX relies on an empirically determined library of rotamers and only considers a frozen local neighborhood of the mutated residue, ignoring the rest of the protein.
  – PROPKA determines pKa values based on a single structure, potentially missing groups that are solvent exposed in certain conformational states.
  – PDB2PQR operates in a purely binary way, assigning discrete protonation states of titratable groups based on a single comparison of the pKa value with the solvent pH, rather than accounting for the equilibrium ratio of protonation.
  – The finite difference solver of APBS makes several assumptions that all have their own drawbacks. Of these assumptions, one of the most prominent is the representation of the solvent as a continuum, rather than a medium composed of individual atoms. This ignores specific interactions of the solvent and ions with the solute (*i*.*e*., the protein), which are likely to play an important role in downstream assayed properties. Further reading on caveats and potential errors of APBS can be found in the official documentation (26).
- **CNN performance may be suboptimal**. There may also be limitations in the feature extraction ability of the 3D CNN. This neural network is relatively small, consisting of only four convolutional layers. In our efforts in hyperparameter tuning, we found that adding additional layers and increasing the number of nodes in the network was not beneficial to performance. However, it is still possible that the limited size of our network impairs the predictive capacity of our model, as the relationship between electrostatics and downstream assayed properties may still be complex.
- **Benchmarking data may be non-representative**. Lastly, benchmarking on ProteinGym datasets may not optimally demonstrate the potential of ChargeNet, as these datasets do not represent typical protein engineering objectives. Protein engineering datasets often measure non-native function – *i*.*e*., protein properties not selected for by evolutionary pressures – and therefore such datasets may be less amenable to models operating solely on evolutionary information. Within this domain, ChargeNet could offer an advantage, as it relies exclusively on physically grounded representations. A larger presence of protein engineering datasets in benchmarking suites could help differentiate the utility of various model archetypes and offer insights into the relationship between protein representations and modeling tasks.

### 3.2 Future directions

Due to computational constraints, a large portion of the hyperparameter space of ChargeNet has not yet been explored. This space is large, as each component of the pipeline is parametrized. The physics module can not leverage GPU computation, and the pipeline is inherently sequential, making it computationally expensive to search the hyperparameter space. For these reasons, we have primarily focused on finding an optimal 3D CNN architecture and have mostly avoided tuning parameters of the physics module. The only exception to this is finding an optimal resolution – *i*.*e*., the number of voxels in the 3D grids – which did not diminish performance when set at the lowest setting allowed by APBS.

Future work could explore different choices in integrating the components that make up the pipeline. One is to implement ChargeNet as a fully differentiable end-to-end model, replacing FoldX with a structure prediction model, and extending this with a differentiable electrostatics module. Doing so would address three problems at once: It could reduce compounding errors as there is now a gradient defined over them, allowing for direct minimization of this effect. It could also shrink the hyperparameter space by allowing the parameters that would normally be pre-configured in the physics module to become trainable, and it would enable full use of GPU computation, facilitating training and more efficient experimentation.

In section 3.1, we pointed out that increasing model complexity did not benefit model performance. On the other hand, we also noted that the final architecture of the 3D CNN is small and may not be expressive enough. One explanation for these observations may simply be that are representations are not accurate enough, but another could be that our method requires a larger amount of data. In addition to simply increasing the size of VEP datasets, an interesting option to explore could be to pretrain a model on electrostatic representations of a diverse set of proteins spanning many families. This could be done in a self-supervised way, *e*.*g*., by predicting precomputed electrostatic quantities from sequence, or in a supervised way by predicting assayed properties of a diverse set of proteins from their electrostatics. The Meltome atlas is an example of a dataset that would fit the latter training method (27).

Another component of our work that we feel is underexplored is the ensembling experiment. Though we have relied on a well established method of combining predictions, we may not have chosen the most effective method for doing so. Perhaps the weights of the ensemble should depend on the individual predictions of its members, and/or on the independent variable – *i*.*e*., the sequences. The stacking ensemble that we created does not account for this; it finds a fixed set of weights for a given dataset. Thus, predictions may not be optimally combined. Using both the model predictions and sequences as inputs to the meta learner may prove to be a more efficient way of complementing evolutionary models with ChargeNet.

### 3.3 Conclusion

We proposed, implemented, and extensively tested a novel method of physics-based VEP. Despite not identifying a use case for ChargeNet, we demonstrated that electrostatic representations have significant predictive capabilities. By documenting our method and experiments in detail, we hope to guide scientists exploring similar methods to focus on different ideas of using electrostatic representations for VEP, and to contribute to the discourse in the field to advance the understanding of the use of physics-based representations in machine learning for structural biology.

## 4. Methods

### 4.1 Data curation

We collected all datasets included in this work from the “DMS Substitutions” benchmark from the ProteinGym repository (1). We selected datasets only if they matched all of the following criteria:

1. There exists a structure with a sequence matching the reference (*i*.*e*., unmutated) sequence of the dataset.
2. The region spanning the first and last positions of mutations in the dataset is entirely resolved in the reference structure.
3. The reference structure and variant sequences do not contain non-canonical amino acids.

This resulted in a total of 22 datasets, five of which could not properly be parsed by PDB2PQR, and were therefore excluded from the analysis: A0A1I9GEU1_NEIME_Kennouche_2019, CALM1_HUMAN_Weile_2017, MYO3_YEAST_Tsuboyama_2023_2BTT, PIN1_HUMAN_Tsuboyama_2023_1I6C, and R1AB_SARS2_Flynn_2022. The final selection of datasets is listed in Table S7.

### 4.2 The PIME CV strategy

The PIME (position inclusive mutation exclusive) CV strategy aims to create folds such that a given position in the sequence is mutated both in the training data and in the test data. The only circumstance under which this is not possible is when the number of variants that contain a mutation at a given position is lower than the number of folds. The PIME CV strategy iterates over each position that is mutated in the dataset, then gathers all variants that have a mutation at that position, and subsequently assigns folds to those variants in a random uniform manner.

### 4.3 The physics module

#### FoldX

We obtained structures of protein variants by mutating the structure corresponding to the reference sequence using the BuildModel command of FoldX 5.0 (28). We used the BuildModel command with default configuration and without changing any of the general parameters.

**PDB2PQR** We prepared mutated structures for continuum electrostatics calculations using PDB2PQR version 3.6.2 (17)(29). We set the force field parameter to AMBER using the `ff` parameter and used the `with-ph` parameter to set the pH in the experiments described in 2.6.

**APBS** We carried out continuum electrostatics computations using APBS version 3.4.1 (29). The configuration of the input file is presented in Listing 1. We used the mg-auto solver of APBS, which implements the finite difference method to compute electrostatics.

The resolution of the computed electrostatics is specified by the `dime` parameter, and this value represents the length of a voxel edge in Å. To illustrate this with an example, a voxel grid with dimensions *d* = (10, 20, 20) at resolution 1.5 would span a region of 15Å × 30Å × 30Å. We set all other parameters to recommended values for large biomolecules, as per the documentation of APBS (30). We used the recommended parameterization without further optimization due to lack of domain-specific expertise and instead relied on established routines for protein electrostatics analysis. We set the `ion` fields to values representing those of typical physiological conditions (31). Lastly, the `write` fields instruct APBS to output the charge distribution, the electrostatic potential and the solvent charge density in *e*_*c*_, *k*_*b*_*T*/*e*_*c*_ and *e*_*c*_*M* respectively, where *e*_*c*_ is the elementary charge, *k*_*b*_ is the Boltzmann constant, *T* is temperature and *M* is molarity.

Listing 1: APBS input file configuration. CENTER, FINE_GRID_LENGTH, and COARSE_GRID_LENGTH are determined by the psize module of PDB2PQR with the reference structure as input, and are therefore dependent on the dataset. PQR_FILE is the PQR file created by PDB2PQR of the variant for which electrostatics calculations are done.

~~~
read
 mol pqr PQR_FILE
end
elec
 mg-auto
 mol 1
 fgcent CENTER
 cgcent CENTER
 fglen FINE_GRID_LENGTH
 cglen COARSE_GRID_LENGTH
 dime 1.5
 lpbe
 bcfl sdh
 pdie 2.0
 sdie 78.0
 chgm spl2
 srfm smol
 swin 0.3
 temp 298
 sdens 10.0
 calcenergy no
 calcforce no srad 1.4
 ion charge +1 conc 0.15 radius 2.0
 ion charge −1 conc 0.15 radius 1.8
 write charge OUTPUT_DIR_CHARGE
 write pot OUTPUT_DIR_POT
 write qdens OUTPUT_DIR_DENS
end
quit
~~~

### 4.4 3D CNN

**Architecture** We implemented the 3D CNN in Pytorch version 1.13.1. It follows a ResNet architecture with just a single residual block, visualized in Figure 1d (16). We set the padding of all convolutional layers to `“same”` to preserve the dimensionality of the feature maps to allow for elementwise summation between residual connections. We set the dropout ratio to 0.3.

**Hyperparameter tuning**

We tuned ChargeNet’s hyperparameters using a separate private dataset with Bayesian optimization provided by Sagemaker version 2.117.0. The Bayesian optimization procedure sampled a total of 100 hyperparameter configurations from the search space. The hyperparameter domains as well as the final configuration are specified in Table 4.

**Table 4.** The ChargeNet hyperparameter domains that are searched by Bayesian optimization.

| Parameter | Search domain | Final value |
| --- | --- | --- |
| Convolutional layer output channels | $\{4, 5, \dots, 32\}$ | 13 |
| CNN residual blocks | $\{1, 2, \dots, 5\}$ | 1 |
| Convolutional kernel edge length | $\{3, 4, \dots, 9\}^1$ | 3 |
| Max pooling window edge length | $\{3, 4, \dots, 9\}^1$ | 3 |
| Loss function | $\{\text{MSE}, \text{MAE}, \text{Huber}, \text{Spearman}\}^2$ | MSE |
| Batch size | $\{8, 9, \dots, 32\}$ | 20 |
| Optimizer | $\{\text{Adam}, \text{Adamw}\}$ | Adamw |
| Optimizer learning rate | $[10^{-5}, 10^{-3}]$ | $10^{-4}$ |
| Optimizer weight decay | $[10^{-6}, 10^{-1}]$ | $5 \times 10^{-6}$ |

<sup>1</sup> Edge length corresponds to all three dimensions of the kernel and pooling window. This means that an edge length of 3 corresponds to a kernel or pooling window size of (3, 3, 3).
<sup>2</sup> We tested a differentiable Spearman loss function using the torchmetrics library (32).

#### Training

After finding the best set of hyperparameters among those identified by the Bayesian optimization, we trained the 3D CNN with a mean squared error loss for 300 epochs with a patience of 20 as an early stopping criterion. This early stopping criterion monitored the loss value on a validation set, obtained by splitting off one of the folds from the training data. We used a batch size of 20 and Pytorch’s `adamw` optimizer with a learning rate of 10^−4^ and a weight decay of 5 *×* 10^−6^. We trained all models on either the ml.g5 or ml.g4dn instances of AWS Sagemaker.

### 4.5 Dipole regressor prediction heads

We implemented the OLS and POLS prediction heads of the dipole regressor in scikit-learn version 1.5.2 (33) using the `LinearRegression` and `PolynomialOLS` classes, respectively. We set the `degree` parameter of the POLS model to 2.

### 4.6 Evolutionary models

We chose two representative evolutionary models: the augmented Potts model and the RITA regressor. These models are described in depth by van der Flier et al. (22). In short, the augmented Potts model uses GREMLIN_CPP v1.0 to fit a Potts model to a multiple sequence alignment (MSA) of the reference sequence. The sequence energy of each variant is then calculated from the field and coupling parameters of the Potts model, and subsequently concatenated with a one-hot encoding vector of the sequence. These feature vectors are used to fit a ridge regressor model with a regularization parameter, *α*, of 0.1.

The RITA regressor uses the RITA_xl protein language model to obtain mean pooled hidden states across the sequence dimensions of protein variants (34). RITA_xl produces hidden states in both directions: N to C and C to N. To combine the hidden states from both directions, we embed variant sequences by concatenating both mean pooled hidden states, resulting in feature vectors of length 4,096. Like the augmented Potts model, the RITA regressor contains a ridge regression prediction head with an *α* of 0.1. We implemented ridge regression models using scikit-learn 1.5.2 (33).

### 4.7 Synthetic pooled solvent charge density prediction

We formulated a prediction task that should be trivial for ChargeNet to solve as a control experiment to validate the implementation of the pipeline. We generated synthetic training labels for dataset HECD1_HUMAN_Tsuboyama_2023_3DKM from the ProteinGym that correspond to the average solvent charge density calculated by the physics module in a predefined region of the protein. Since these features are derived from the physics module with an identical configuration as the physics module in ChargeNet used for prediction, we expect ChargeNet to perform well on this task. To ensure that the task itself is not trivial, we separate the dataset into training, validation, and testing based on the positions that are mutated, such that there is no overlap in positions between the sets.

#### Training target generation

We begin by generating solvent charge densities of all variants in the dataset with the physics module using the configuration as described in Methods 4.3. We then pick a pooling window within which the average solvent charge density will be computed for all variants. We opted for a window of (7, 7, 7) voxels, because the corresponding size of the volume is be 10.5Å *×*10.5Å*×* 10.5Å when running APBS with a resolution of 1.5Å. A volume of this size roughly matches that of many small molecules and is therefore biologically meaningful. We perform an average pooling operation on all the computed solvent charge densities using Pytorch’s `AvgPool3d` operation with a stride of 1. From these pooled voxel representations of solvent charge density, we pick the voxel that displays the highest pooled solvent charge density variance across all variants in the dataset to ensure maximum separation in target space. For a given variant, the value at this voxel is set as the training target. Distributions of the training targets by split are plotted in Figure S1a.

#### Evaluation

We create three sets of positions without overlap, and assign each of them to a split. Specifically, the training set contains variants with mutations at residues {R15, D21, W24, T36, D51, R59, E63, K65, D67}, the validation set at residues {R18, D26, D28} and the test set at residues {Q27, W50, N56, Y58}, with position indexing corresponding to the reference sequence as found in the ProteinGym. Figure S1b illustrates where these residues are located in the structure of the reference, *i*.*e*., unmutated protein. Predictors are evaluated by training on 10 sets that are subsampled down to 60% of the entire dataset, with the subsampling applied to each split individually to ensure that relative split sizes remain similar. The RITA regressor and the augmented Potts model had the validation data assigned to the training data, whereas the validation data for ChargeNet was used as an early stopping criterion. We assessed performance by Spearman correlation of the test predictions with the test labels for all models, which are presented in Table S8.

### 4.8 Ensembling

In an attempt to improve model predictions, we opted for an ensembling technique referred to as stacking (23). A stacking ensemble consists of base learners – *i*.*e*., the models we aim to combine – and a meta learner that fits base learner predictions to ground truth values, which in our case is a ridge regression model. By choosing a linear regression model as our meta learner, our ensemble predictions can be interpreted as a weighted average of base learner predictions offset by a bias term. We describe the nested CV procedure for the stacked ensemble in Algorithm 1.

We repeated the stacking procedure for all combinations of models, including single models. Though this may seem redundant, we did so to create a fair comparison of standalone models with meta learner models that have identical training routines – *i*.*e*., are exposed to identical data points in the nested CV.

#### Algorithm 1

Nested cross-validation of a stacked ensemble.

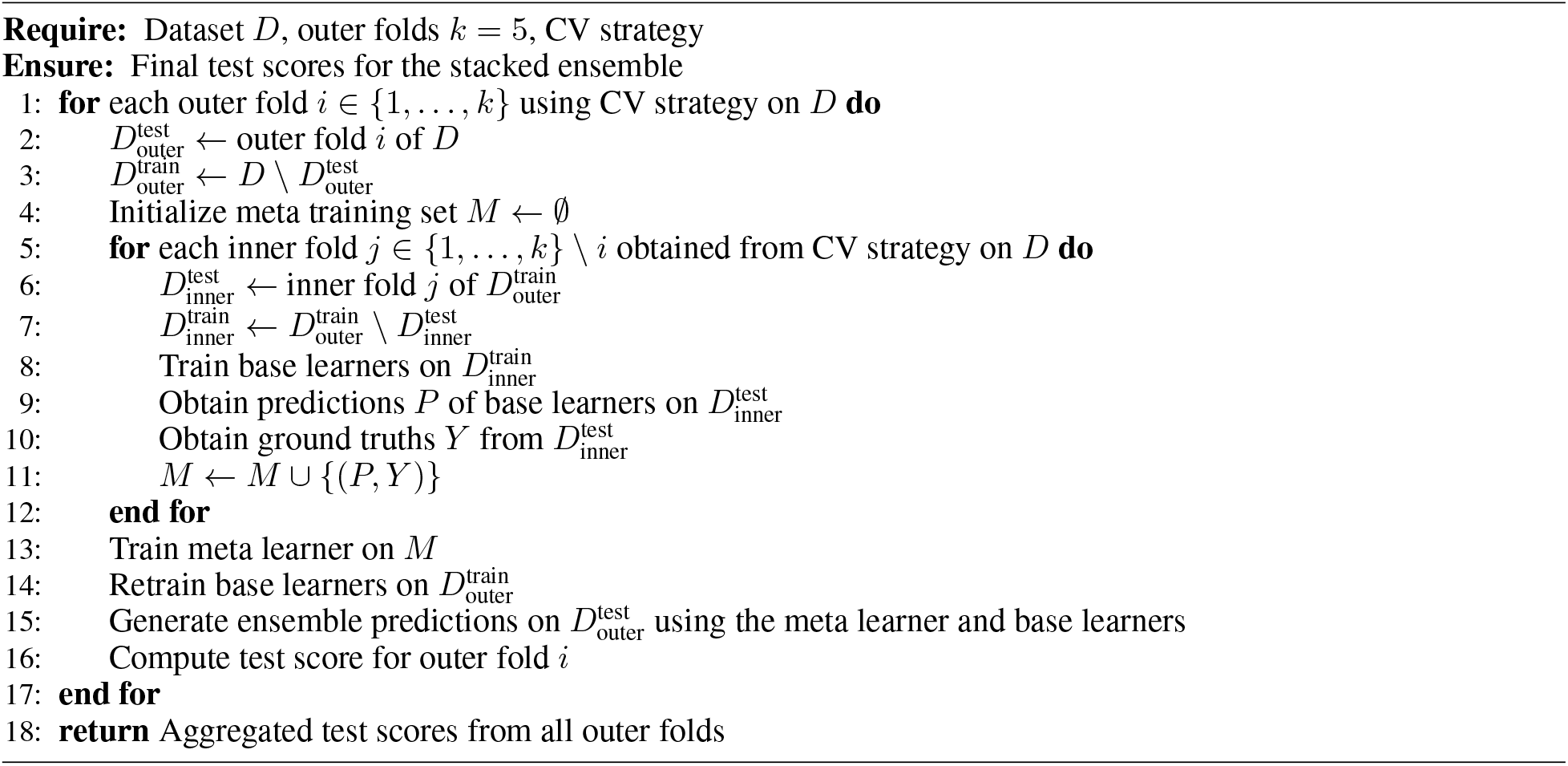

## Acknowledgments

We express our gratitude to Luis Cascão-Pereira and David Estell for sharing their expertise and insights about the role of electrostatics in protein engineering screening data, which ultimately drove us to develop ChargeNet.

## Data availability

We obtained all datasets used in this analysis from the ProteinGym (1), except the *α*-amylase dataset, which was obtained from the pilot edition of the Protein Engineering Tournament github repository: https://github.com/the-protein-engineering-tournament/pet-pilot-2023 (24).

## Code availability

Code is made available at https://github.com/florisvdf/chargenet.

## Declaration of generative AI and AI-assisted technologies in the writing process

During the writing of the article, authors used various GPT models developed by OpenAI to assist with grammar, flow, and overall refinement of the text. After using these tools, the authors carefully reviewed and edited the generated content as needed and take full responsibility for the content of the published article.

## A Supplementary Materials

**Figure S1.**
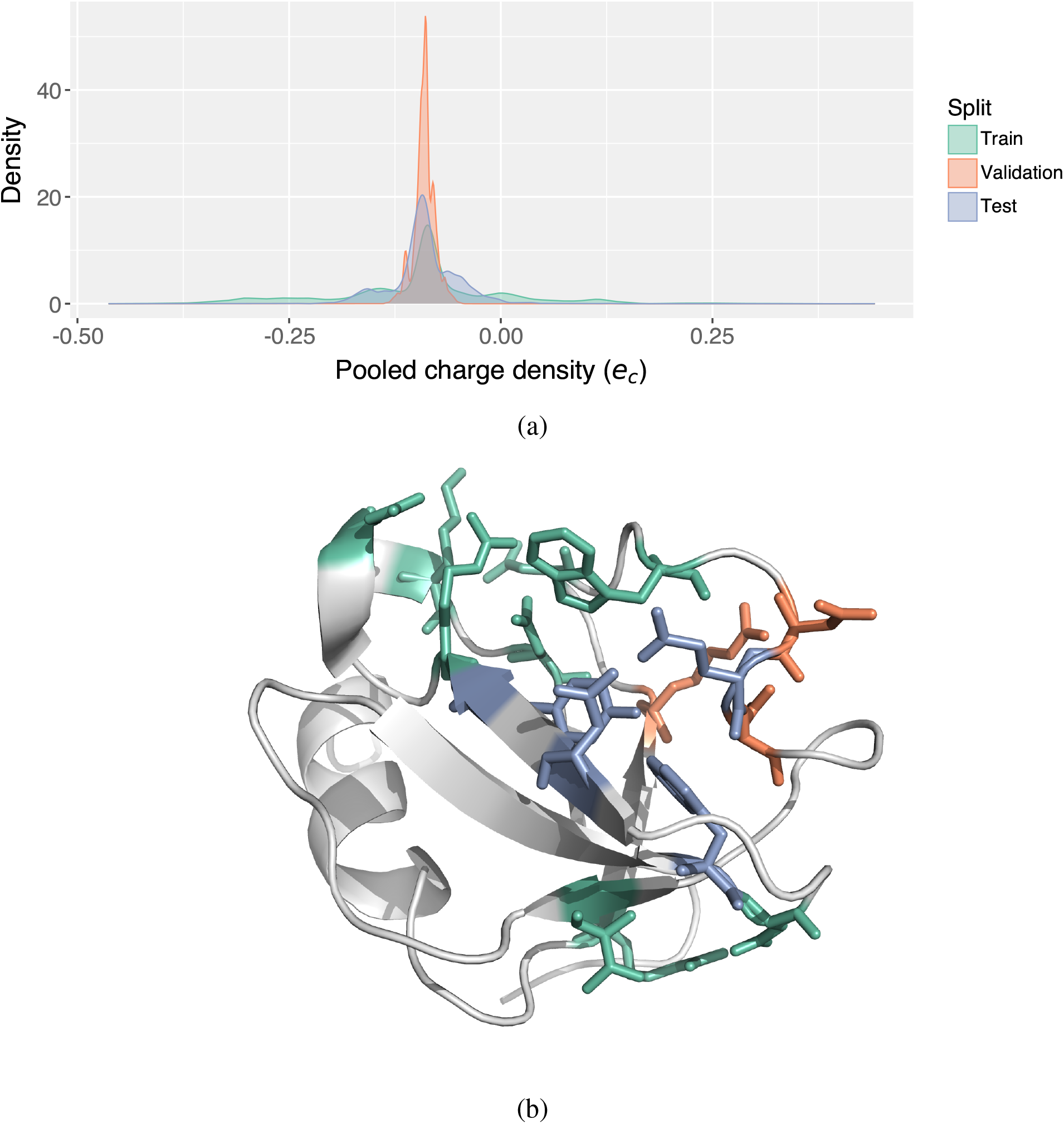
(a) Distribution of the average pooled computed solvent charge density by data split plotted as kernel density estimations. (b) The reference structure of the HECD1_HUMAN_Tsuboyama_2023_3DKM dataset (PDB ID: pdb_00003dkm). The structure presented to ChargeNet and used to compute pooled solvent charge densities had its leading residues MHHHHHHSSGRE and trailing residues GYDPD removed, since they do not appear in the reference sequence. Residues are colored by the split in which they are mutated, with the coloring matching the legend in (a). Concretely, residues mutated in the training, validation and test set are: {R15, D21, W24, T36, D51, R59, E63, K65, D67}, {R18, D26, D28} and {Q27, W50, N56, Y58}, respectively, with position indexing corresponding to the reference sequence as found in the ProteinGym.

**Table S1.**
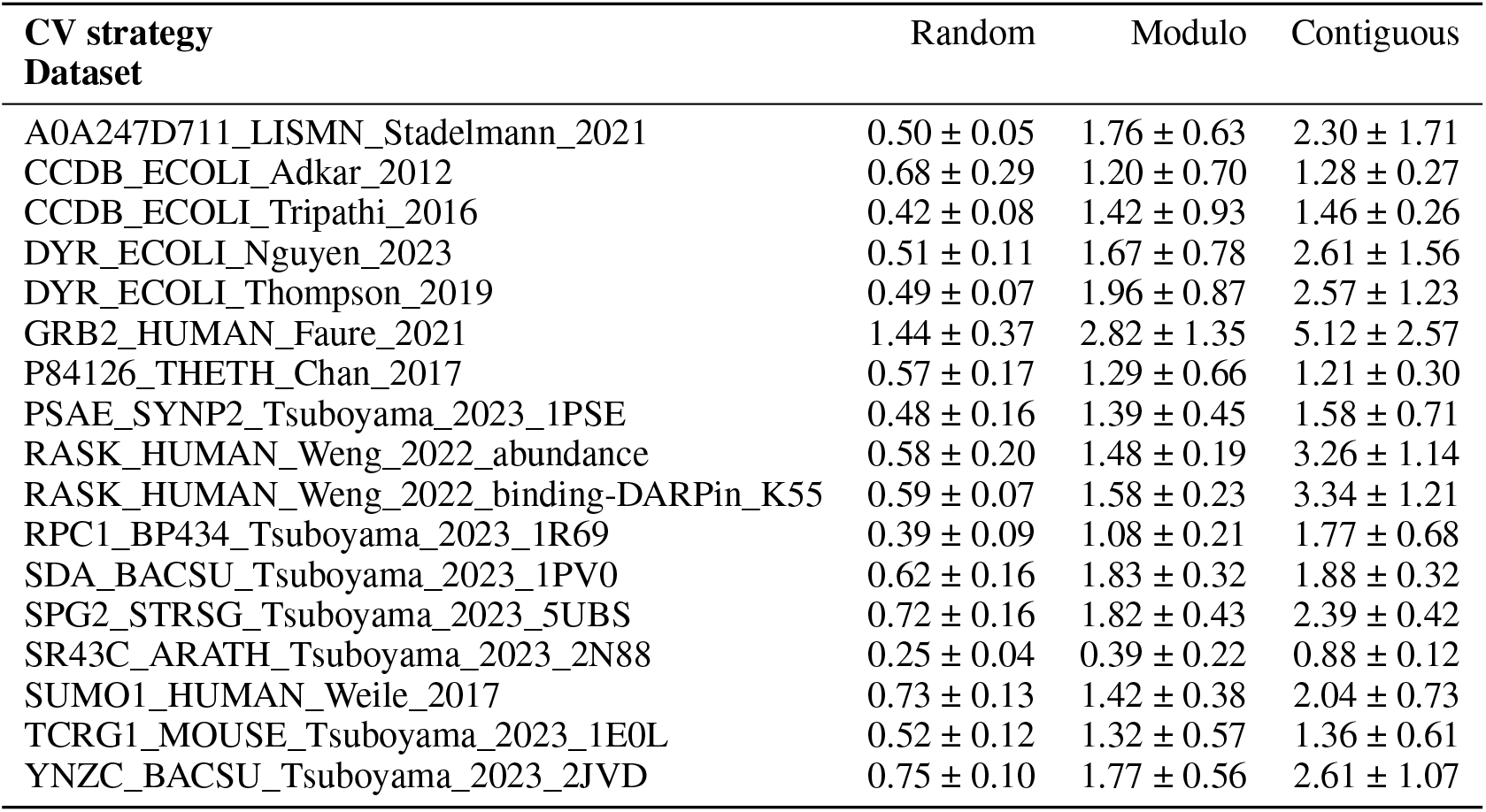
Averages and standard deviations of intra-fold Wasserstein distances of the dipole moment magnitudes for all three splitting strategies for the curated protein gym datasets. Higher values indicate a lower degree of similarity. For univariate distributions, the Wasserstein distance is the area under the absolute difference between the two cumulative density functions. For a given split and dataset, Wasserstein distances are computed between each test fold and the remaining folds across all five folds.

| CV strategy<br>Dataset | Random | Modulo | Contiguous |
| --- | --- | --- | --- |
| A0A247D711_LISMN_Stadelmann_2021 | $0.50 \pm 0.05$ | $1.76 \pm 0.63$ | $2.30 \pm 1.71$ |
| CCDB_ECOLI_Adkar_2012 | $0.68 \pm 0.29$ | $1.20 \pm 0.70$ | $1.28 \pm 0.27$ |
| CCDB_ECOLI_Tripathi_2016 | $0.42 \pm 0.08$ | $1.42 \pm 0.93$ | $1.46 \pm 0.26$ |
| DYR_ECOLI_Nguyen_2023 | $0.51 \pm 0.11$ | $1.67 \pm 0.78$ | $2.61 \pm 1.56$ |
| DYR_ECOLI_Thompson_2019 | $0.49 \pm 0.07$ | $1.96 \pm 0.87$ | $2.57 \pm 1.23$ |
| GRB2_HUMAN_Faure_2021 | $1.44 \pm 0.37$ | $2.82 \pm 1.35$ | $5.12 \pm 2.57$ |
| P84126_THETH_Chan_2017 | $0.57 \pm 0.17$ | $1.29 \pm 0.66$ | $1.21 \pm 0.30$ |
| PSAE_SYNP2_Tsuboyama_2023_1PSE | $0.48 \pm 0.16$ | $1.39 \pm 0.45$ | $1.58 \pm 0.71$ |
| RASK_HUMAN_Weng_2022_abundance | $0.58 \pm 0.20$ | $1.48 \pm 0.19$ | $3.26 \pm 1.14$ |
| RASK_HUMAN_Weng_2022_binding-DARPin_K55 | $0.59 \pm 0.07$ | $1.58 \pm 0.23$ | $3.34 \pm 1.21$ |
| RPC1_BP434_Tsuboyama_2023_1R69 | $0.39 \pm 0.09$ | $1.08 \pm 0.21$ | $1.77 \pm 0.68$ |
| SDA_BACSU_Tsuboyama_2023_1PV0 | $0.62 \pm 0.16$ | $1.83 \pm 0.32$ | $1.88 \pm 0.32$ |
| SPG2_STRSG_Tsuboyama_2023_5UBS | $0.72 \pm 0.16$ | $1.82 \pm 0.43$ | $2.39 \pm 0.42$ |
| SR43C_ARATH_Tsuboyama_2023_2N88 | $0.25 \pm 0.04$ | $0.39 \pm 0.22$ | $0.88 \pm 0.12$ |
| SUMO1_HUMAN>Weile_2017 | $0.73 \pm 0.13$ | $1.42 \pm 0.38$ | $2.04 \pm 0.73$ |
| TCRG1_MOUSE_Tsuboyama_2023_1E0L | $0.52 \pm 0.12$ | $1.32 \pm 0.57$ | $1.36 \pm 0.61$ |
| YNZC_BACSU_Tsuboyama_2023_2JVD | $0.75 \pm 0.10$ | $1.77 \pm 0.56$ | $2.61 \pm 1.07$ |

**Table S2.** Averages and standard deviations of Spearman correlation coefficient scores of the dipole regressor for all datasets, CV strategies, and regressor head. Model name abbreviations are: AP: augmented Potts model, RR: RITA regressor, CN: ChargeNet. Dataset indices correspond to the order in Table S1.

| CV strategy<br>Regressor head<br>Dataset index | Random<br>OLS | POLS | Modulo<br>OLS | POLS | Contiguous<br>OLS | POLS |
| --- | --- | --- | --- | --- | --- | --- |
| 0 | 0.05 ± 0.06 | 0.11 ± 0.04 | -0.16 ± 0.14 | -0.09 ± 0.14 | -0.04 ± 0.10 | -0.09 ± 0.16 |
| 1 | 0.15 ± 0.11 | 0.38 ± 0.06 | 0.06 ± 0.19 | 0.31 ± 0.17 | 0.12 ± 0.10 | 0.33 ± 0.10 |
| 2 | 0.11 ± 0.06 | 0.17 ± 0.05 | 0.07 ± 0.08 | 0.12 ± 0.06 | 0.06 ± 0.08 | 0.06 ± 0.05 |
| 3 | 0.07 ± 0.03 | 0.16 ± 0.02 | -0.06 ± 0.13 | 0.06 ± 0.10 | -0.04 ± 0.17 | 0.05 ± 0.20 |
| 4 | 0.07 ± 0.04 | 0.20 ± 0.06 | -0.11 ± 0.07 | 0.08 ± 0.09 | 0.04 ± 0.10 | 0.07 ± 0.10 |
| 5 | 0.06 ± 0.07 | 0.11 ± 0.11 | -0.03 ± 0.14 | 0.04 ± 0.06 | 0.00 ± 0.19 | 0.02 ± 0.19 |
| 6 | 0.02 ± 0.10 | 0.14 ± 0.03 | -0.07 ± 0.12 | 0.06 ± 0.05 | -0.06 ± 0.11 | -0.09 ± 0.07 |
| 7 | 0.04 ± 0.07 | 0.03 ± 0.06 | -0.09 ± 0.16 | -0.11 ± 0.15 | -0.04 ± 0.12 | -0.07 ± 0.19 |
| 8 | 0.09 ± 0.05 | 0.13 ± 0.06 | 0.05 ± 0.03 | 0.07 ± 0.07 | 0.05 ± 0.06 | 0.08 ± 0.05 |
| 9 | -0.03 ± 0.05 | 0.20 ± 0.04 | -0.08 ± 0.06 | 0.19 ± 0.10 | -0.14 ± 0.13 | 0.05 ± 0.11 |
| 10 | 0.14 ± 0.04 | 0.17 ± 0.05 | 0.11 ± 0.14 | 0.07 ± 0.11 | -0.04 ± 0.25 | 0.05 ± 0.24 |
| 11 | 0.11 ± 0.03 | 0.24 ± 0.07 | -0.00 ± 0.36 | -0.15 ± 0.13 | 0.02 ± 0.20 | -0.01 ± 0.17 |
| 12 | -0.03 ± 0.08 | 0.03 ± 0.08 | -0.23 ± 0.05 | -0.12 ± 0.12 | -0.19 ± 0.12 | -0.23 ± 0.15 |
| 13 | 0.13 ± 0.05 | 0.15 ± 0.04 | 0.11 ± 0.17 | 0.07 ± 0.16 | -0.02 ± 0.09 | -0.08 ± 0.12 |
| 14 | 0.13 ± 0.04 | 0.18 ± 0.04 | 0.09 ± 0.07 | 0.13 ± 0.09 | 0.07 ± 0.07 | 0.02 ± 0.12 |
| 15 | 0.09 ± 0.07 | 0.13 ± 0.07 | 0.02 ± 0.21 | 0.01 ± 0.19 | 0.06 ± 0.05 | 0.10 ± 0.13 |
| 16 | 0.27 ± 0.13 | 0.37 ± 0.08 | 0.13 ± 0.06 | 0.30 ± 0.17 | 0.09 ± 0.22 | 0.32 ± 0.11 |

**Table S3.** Averages and standard deviations of Spearman correlation coefficient scores of ChargeNet, the augmented Potts model, and the RITA regressor for all curated ProteinGym datasets over all five folds using the random CV strategy.

| Model<br>Dataset | ChargeNet | Augmented Potts model | RITA regressor |
| --- | --- | --- | --- |
| A0A247D711_LISMN_Stadelmann_2021 | 0.52 ± 0.05 | **0.61 ± 0.07** | 0.59 ± 0.05 |
| CCDB_ECOLI_Adkar_2012 | 0.80 ± 0.03 | **0.97 ± 0.00** | 0.81 ± 0.02 |
| CCDB_ECOLI_Tripathi_2016 | 0.51 ± 0.04 | **0.60 ± 0.05** | 0.59 ± 0.05 |
| DYR_ECOLI_Nguyen_2023 | 0.51 ± 0.06 | **0.62 ± 0.04** | 0.61 ± 0.06 |
| DYR_ECOLI_Thompson_2019 | 0.51 ± 0.03 | **0.69 ± 0.02** | 0.65 ± 0.03 |
| GRB2_HUMAN_Faure_2021 | 0.64 ± 0.02 | 0.80 ± 0.02 | **0.81 ± 0.02** |
| P84126_THETH_Chan_2017 | 0.55 ± 0.07 | **0.78 ± 0.02** | 0.77 ± 0.02 |
| PSAE_SYNP2_Tsuboyama_2023_1PSE | 0.67 ± 0.05 | 0.74 ± 0.04 | **0.81 ± 0.04** |
| RASK_HUMAN_Weng_2022_abundance | 0.54 ± 0.03 | 0.67 ± 0.03 | **0.78 ± 0.01** |
| RASK_HUMAN_Weng_2022_binding-DARPin_K55 | 0.60 ± 0.04 | 0.75 ± 0.01 | **0.82 ± 0.01** |
| RPC1_BP434_Tsuboyama_2023_1R69 | 0.66 ± 0.06 | 0.82 ± 0.02 | **0.91 ± 0.01** |
| SDA_BACSU_Tsuboyama_2023_1PV0 | 0.61 ± 0.04 | 0.75 ± 0.03 | **0.83 ± 0.03** |
| SPG2_STRSG_Tsuboyama_2023_5UBS | 0.60 ± 0.03 | 0.65 ± 0.03 | **0.84 ± 0.01** |
| SR43C_ARATH_Tsuboyama_2023_2N88 | 0.52 ± 0.08 | 0.80 ± 0.03 | **0.83 ± 0.03** |
| SUMO1_HUMAN>Weile_2017 | 0.46 ± 0.03 | **0.63 ± 0.02** | 0.60 ± 0.04 |
| TCRG1_MOUSE_Tsuboyama_2023_1E0L | 0.71 ± 0.05 | 0.80 ± 0.01 | **0.91 ± 0.01** |
| YNZC_BACSU_Tsuboyama_2023_2JVD | 0.63 ± 0.06 | 0.72 ± 0.05 | **0.88 ± 0.03** |

**Table S4.**
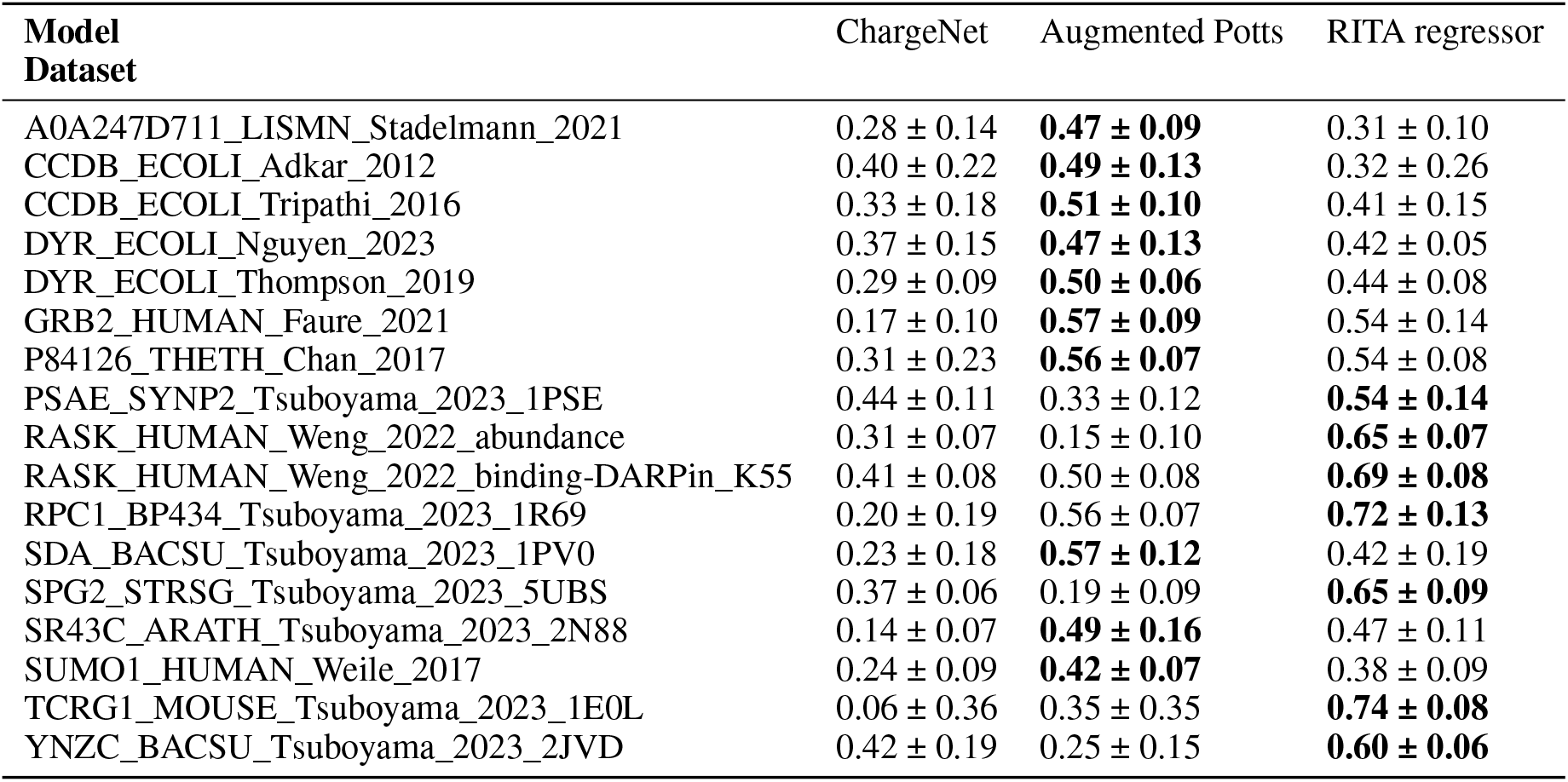
Averages and standard deviations of Spearman correlation coefficient scores of ChargeNet, the augmented Potts model, and the RITA regressor for all curated ProteinGym datasets over all five folds using the modulo CV strategy.

**Table S5.** Averages and standard deviations of Spearman correlation coefficient scores of ChargeNet, the augmented Potts model, and the RITA regressor for all curated ProteinGym datasets over all five folds using the contiguous CV strategy.

| Model<br>Dataset | ChargeNet | Augmented Potts model | RITA regressor |
| --- | --- | --- | --- |
| A0A247D711_LISMN_Stadelmann_2021 | 0.28 ± 0.14 | **0.41 ± 0.16** | 0.23 ± 0.10 |
| CCDB_ECOLI_Adkar_2012 | 0.22 ± 0.13 | **0.46 ± 0.11** | 0.12 ± 0.22 |
| CCDB_ECOLI_Tripathi_2016 | 0.22 ± 0.11 | **0.48 ± 0.12** | 0.15 ± 0.09 |
| DYR_ECOLI_Nguyen_2023 | 0.33 ± 0.09 | **0.45 ± 0.17** | 0.40 ± 0.19 |
| DYR_ECOLI_Thompson_2019 | 0.12 ± 0.10 | **0.50 ± 0.06** | 0.36 ± 0.09 |
| GRB2_HUMAN_Faure_2021 | 0.27 ± 0.18 | **0.63 ± 0.08** | 0.52 ± 0.09 |
| P84126_THETH_Chan_2017 | 0.27 ± 0.09 | **0.62 ± 0.08** | 0.56 ± 0.19 |
| PSAE_SYNP2_Tsuboyama_2023_1PSE | 0.32 ± 0.10 | 0.34 ± 0.11 | **0.54 ± 0.12** |
| RASK_HUMAN_Weng_2022_abundance | 0.21 ± 0.14 | 0.15 ± 0.03 | **0.61 ± 0.07** |
| RASK_HUMAN_Weng_2022_binding-DARPin_K55 | 0.37 ± 0.13 | 0.49 ± 0.07 | **0.67 ± 0.06** |
| RPC1_BP434_Tsuboyama_2023_1R69 | 0.26 ± 0.17 | 0.62 ± 0.08 | **0.70 ± 0.14** |
| SDA_BACSU_Tsuboyama_2023_1PV0 | 0.36 ± 0.11 | **0.62 ± 0.07** | 0.45 ± 0.16 |
| SPG2_STRSG_Tsuboyama_2023_5UBS | 0.25 ± 0.09 | 0.16 ± 0.17 | **0.63 ± 0.06** |
| SR43C_ARATH_Tsuboyama_2023_2N88 | 0.05 ± 0.12 | 0.32 ± 0.17 | **0.41 ± 0.12** |
| SUMO1_HUMAN>Weile_2017 | 0.10 ± 0.13 | **0.38 ± 0.13** | 0.26 ± 0.15 |
| TCRG1_MOUSE_Tsuboyama_2023_1E0L | 0.23 ± 0.23 | 0.32 ± 0.33 | **0.70 ± 0.08** |
| YNZC_BACSU_Tsuboyama_2023_2JVD | 0.35 ± 0.18 | 0.29 ± 0.15 | **0.50 ± 0.19** |

**Table S6.** Averages and standard deviations of Spearman correlation coefficient scores of ensembles of ChargeNet, the augmented Potts model, the RITA regressor, and standalone models for all curated ProteinGym datasets over all five folds using the modulo CV strategy. Dataset indices correspond to the order in Table S1.

| Ensemble<br>Dataset index | AP | RR | CN | AP + RR | AP + CN | RR + CN | AP + RR + CN |
| --- | --- | --- | --- | --- | --- | --- | --- |
| 0 | **0.47 ± 0.09** | 0.31 ± 0.10 | 0.32 ± 0.16 | 0.44 ± 0.10 | 0.46 ± 0.13 | 0.35 ± 0.13 | 0.46 ± 0.12 |
| 1 | 0.49 ± 0.13 | 0.32 ± 0.26 | 0.43 ± 0.15 | 0.49 ± 0.16 | **0.55 ± 0.10** | 0.44 ± 0.23 | 0.55 ± 0.14 |
| 2 | 0.51 ± 0.10 | 0.41 ± 0.15 | 0.35 ± 0.17 | 0.54 ± 0.10 | 0.53 ± 0.12 | 0.45 ± 0.15 | **0.55 ± 0.11** |
| 3 | 0.47 ± 0.13 | 0.42 ± 0.05 | 0.40 ± 0.11 | 0.49 ± 0.11 | 0.50 ± 0.14 | 0.49 ± 0.10 | **0.52 ± 0.13** |
| 4 | 0.50 ± 0.06 | 0.44 ± 0.08 | 0.28 ± 0.09 | **0.54 ± 0.05** | 0.51 ± 0.07 | 0.46 ± 0.09 | 0.54 ± 0.05 |
| 5 | 0.57 ± 0.09 | 0.54 ± 0.14 | 0.10 ± 0.21 | 0.63 ± 0.10 | 0.58 ± 0.08 | 0.56 ± 0.13 | **0.64 ± 0.09** |
| 6 | 0.56 ± 0.07 | 0.54 ± 0.08 | 0.29 ± 0.14 | **0.60 ± 0.07** | 0.56 ± 0.07 | 0.54 ± 0.08 | 0.60 ± 0.08 |
| 7 | 0.33 ± 0.12 | 0.54 ± 0.14 | 0.46 ± 0.16 | 0.55 ± 0.12 | 0.51 ± 0.16 | 0.58 ± 0.13 | **0.60 ± 0.13** |
| 8 | 0.15 ± 0.10 | 0.65 ± 0.07 | 0.32 ± 0.04 | **0.65 ± 0.07** | 0.28 ± 0.12 | 0.65 ± 0.08 | 0.65 ± 0.08 |
| 9 | 0.50 ± 0.08 | 0.69 ± 0.08 | 0.42 ± 0.08 | **0.69 ± 0.08** | 0.54 ± 0.09 | 0.69 ± 0.09 | 0.69 ± 0.09 |
| 10 | 0.56 ± 0.07 | 0.72 ± 0.13 | 0.19 ± 0.20 | **0.76 ± 0.10** | 0.57 ± 0.09 | 0.72 ± 0.13 | 0.75 ± 0.10 |
| 11 | 0.57 ± 0.12 | 0.42 ± 0.19 | 0.16 ± 0.23 | **0.63 ± 0.09** | 0.55 ± 0.15 | 0.43 ± 0.14 | 0.62 ± 0.10 |
| 12 | 0.19 ± 0.09 | 0.65 ± 0.09 | 0.38 ± 0.06 | **0.66 ± 0.09** | 0.37 ± 0.04 | 0.63 ± 0.10 | 0.64 ± 0.11 |
| 13 | 0.49 ± 0.16 | 0.47 ± 0.11 | 0.02 ± 0.11 | **0.59 ± 0.17** | 0.50 ± 0.16 | 0.48 ± 0.11 | 0.59 ± 0.17 |
| 14 | 0.42 ± 0.07 | 0.38 ± 0.09 | 0.24 ± 0.12 | **0.48 ± 0.07** | 0.43 ± 0.07 | 0.38 ± 0.08 | 0.47 ± 0.06 |
| 15 | 0.35 ± 0.35 | **0.74 ± 0.08** | -0.04 ± 0.19 | 0.69 ± 0.13 | 0.22 ± 0.29 | 0.74 ± 0.08 | 0.69 ± 0.12 |
| 16 | 0.25 ± 0.15 | 0.60 ± 0.06 | 0.38 ± 0.23 | 0.57 ± 0.09 | 0.37 ± 0.22 | **0.60 ± 0.09** | 0.58 ± 0.10 |

**Table S7.**
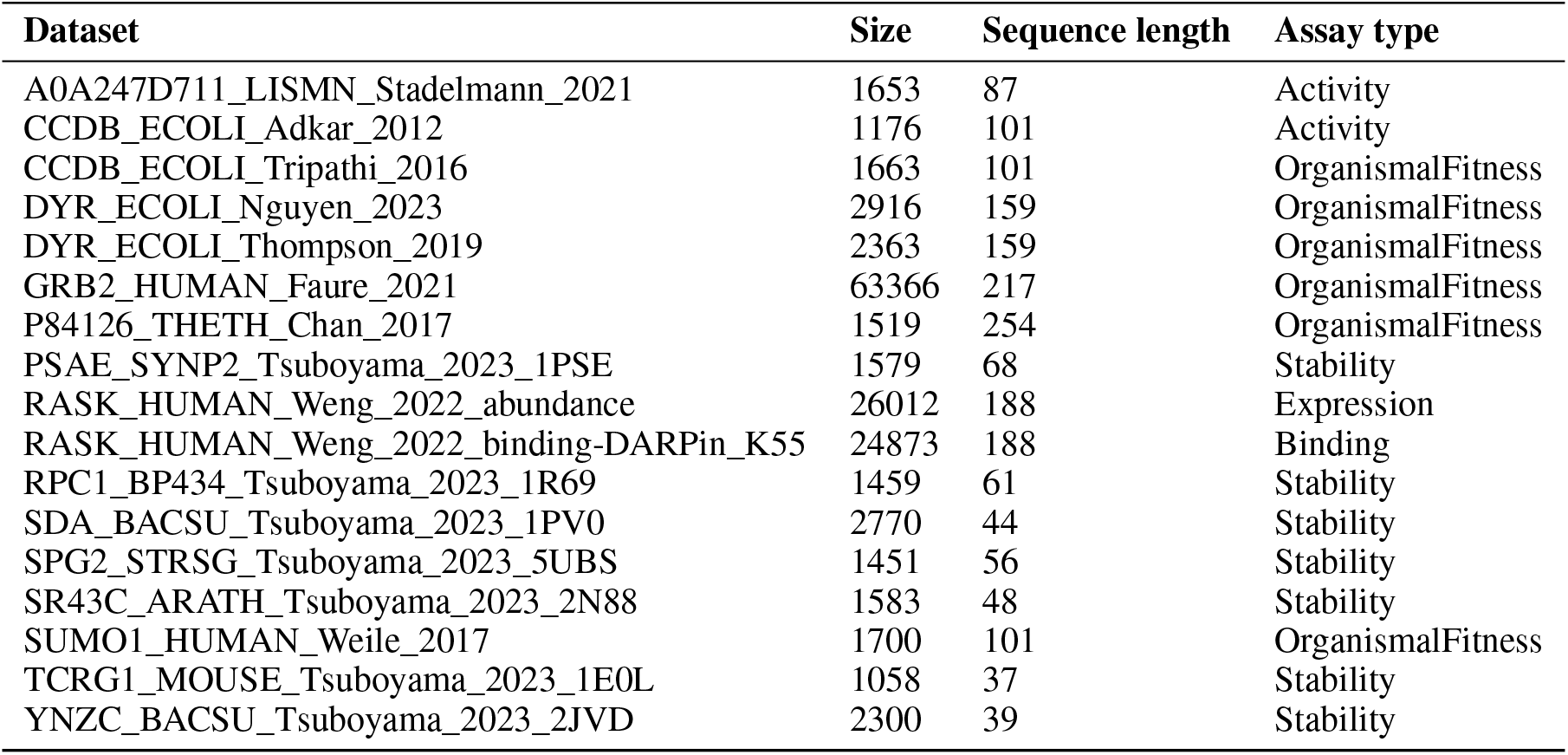
Curated ProteinGym datasets used in our analysis, the size of the datasets, and their assay types.

**Table S8.** Averages and standard deviations of Spearman correlation coefficient scores of ChargeNet, the RITA regressor, and the augmented Potts model on the HECD1_HUMAN_Tsuboyama_2023_3DKM dataset randomly subsampled 10 times at 60%. ChargeNet accurately fits the derived data, indicating that the pipeline behaves as expected and passes the control experiment. In contrast, the augmented Potts model fails to capture the signal, and the RITA regressor shows only marginal predictive performance.

| Augmented Potts model | RITA regressor | ChargeNet |
| --- | --- | --- |
| -0.09 ± 0.02 | 0.26 ± 0.09 | 0.70 ± 0.06 |

